# *Campylobacter jejuni* induces differentiation of human neutrophils to the CD16^hi^/CD62L^lo^ subtype which possess cancer promoting activities

**DOI:** 10.1101/2022.01.28.478176

**Authors:** Carolina G. Dolislager, Sean M. Callahan, Dallas R. Donohoe, Jeremiah G. Johnson

## Abstract

The discovery of neutrophil subtypes has expanded what is known about neutrophil functions, yet there is still much to learn about the role of these subtypes during bacterial infection. We investigated whether *Campylobacter jejuni* induced differentiation of human neutrophils into the hypersegmented, CD16^hi^/CD62L^lo^ subtype. In addition, we investigated whether *C. jejuni*-dependent differentiation of this neutrophil subtype induced cancer promoting activities of human T cells and colonocytes, which were observed in other studies of hypersegmented, CD16^hi^/CD62L^lo^ neutrophils. We found that *C. jejuni* causes a significant shift in human neutrophil populations to the hypersegmented, CD16^hi^/CD62L^lo^ subtype and that those populations exhibit delayed apoptosis, elevated arginase-1 expression, and increased reactive oxygen species production. Furthermore, incubation of *C. jejuni*-infected neutrophils with human T cells resulted in decreased expression of the ζ-chain of the T cell receptor (TCRζ), which was restored upon supplementation with exogenous L-arginine. In addition, incubation of *C. jejuni-*infected neutrophils with human colonocytes resulted in increased HIF-1α stabilization and NF-κB activation in those colonocytes, which may result in the upregulation of pro-tumorigenic genes. This response, coupled with the ability of *C. jejuni-*infected neutrophils to suppress TCRζ expression in T cells, could result in the promotion of colorectal tumorigenesis during infection with *C. jejuni*.

## Introduction

Neutrophils are the most abundant leukocyte in the human body and are often the first immune cells recruited to the site of infection. At this site, neutrophils exert various antimicrobial functions such as phagocytosis, degranulation, reactive oxygen species (ROS) production, and the generation of neutrophil extracellular traps (NETs). Importantly, many of these activities are potent inducers of inflammation and can damage surrounding host tissue [1]. Neutrophils were once thought to be homogeneous, short-lived, and transcriptionally inert, yet much has been learned about their diversity and complex functions in recent years [2]. Neutrophil heterogeneity was first observed in response to *Staphylococcus aureus* infection, with neutrophil subtype differences in cell surface marker and toll-like receptor (TLR) expression, cytokine and chemokine production, and altered ability to activate macrophages [3]. In another study, neutrophil subtypes with antitumorigenic or protumorigenic characteristics were discovered in tumor-associated neutrophil (TAN) populations, which led to the distinction of N1 and N2 TAN subtypes, respectively [4]. Lastly, experimental administration of endotoxin to patients resulted in neutrophil subtypes that are characterized by their nuclear morphology (banded, segmented, or hypersegmented) and expression levels of the cell surface markers CD16 (Fc-gamma III receptor) and CD62L (L-selectin) [5, 6]. Neutrophils expressing high levels of both CD16 and CD62L are typically associated with segmented nuclei with approximately four lobes and are most commonly isolated from the bloodstream under normal conditions [6]. Neutrophils expressing low levels of CD16 and high levels of CD62L present with banded nuclei, tend to be immature, and are released from the bone marrow in instances of trauma and injury [7, 8]. Neutrophils expressing low levels of CD16 and CD62L are often associated with aging and apoptosis [6]. Of particular interest in many studies are neutrophils that highly express CD16, express low levels of CD62L, and possess hypersegmented nuclei. These hypersegmented, CD16^hi^/CD62L^lo^ neutrophils have been observed in response to several conditions such as sterile inflammation (endotoxemia), septic shock, cancer, viral infection with lymphocytic choriomeningitis virus (LCMV), and bacterial infection with *Helicobacter pylori* [5,9,10,11,12]. These neutrophils have been demonstrated to have a proteome distinct from banded and segmented neutrophils and have been observed to differentiate before release into the bloodstream [13]. Once differentiated, CD16^hi^/CD62L^lo^ neutrophils can impact the adaptive immune system through elevated production of arginase-1 and ROS, both of which can reduce expression of the ζ-chain of the T cell receptor (TCRζ), which negatively impacts the ability of T cells to become activated and proliferate [14].

Arginase-1 is an enzyme possessed by granulocytes that converts L-arginine into L-ornithine and urea, which leads to L-arginine depletion. This depletion reduces expression of TCRζ since T cells must import adequate amounts of extracellular L-arginine to properly express TCRζ and proliferate [15, 16]. Moreover, T cells require large amounts of L-arginine during activation [17]. Reactive oxygen species also reduce TCRζ expression, which has been reported to occur via a reduction in mRNA levels and/or direct alteration of proteins without affecting other TCR components like CD3ε (CD3) [18]. To our knowledge, the only bacteria which has been observed to promote CD16^hi^/CD62L^lo^ subtype differentiation is *H. pylori*, where these neutrophils were also shown to have an extended lifespan [12]. In our study, we investigated whether the closely related gastrointestinal pathogen *Campylobacter jejuni* induces similar effects to build the field’s understanding of bacterially-induced differentiation of neutrophil subtypes and their potential downstream effects on the host.

*Campylobacter* spp. are a leading cause of bacterially derived gastroenteritis worldwide, with infection often presenting as a self-limiting, moderate-to-severe inflammatory diarrhea [19, 20]. Recently, asymptomatic carriage of *Campylobacter* spp. has become increasingly appreciated due to its association with the development of environmental enteric dysfunction (EED) in pediatric populations in the developing world [21, 22]. As a result, in cases of *Campylobacter* carriage, the impact of the pathogen on human health may need to be considered similar to *H. pylori*, where infection can persist [23]. Beyond this, there are several similarities that exist between what has been found during infection with *H. pylori* and *C. jejuni*. For example, both *H. pylori* and *C. jejuni* infection result in high levels of neutrophil recruitment and activity, which result in severe inflammation at the site of infection [24,25,26]. Importantly, the chronic neutrophilic response to *H. pylori* infection is believed to result from immune dysregulation and has been suggested to contribute to the development of gastric cancer [27]. Further, *H. pylori* infection results in DNA damage due to the generation of extracellular ROS by recruited neutrophils and the bacterium itself, which contributes to the formation of a protumorigenic environment [28]. Similarly, *C. jejuni* can induce DNA damage through secretion of cytolethal distending toxin (CDT), which enters the nucleus where it exhibits type I DNase activity and catalyzes double-strand breaks [29]. While *H. pylori*-associated gastric cancer has been well established, and despite the similar activities mentioned above and the prevalence of *Campylobacter* spp. infection, much less is known about the relationship between *Campylobacter* spp. and cancer [30,31,32].

One previous study followed Swedish patients who had previously had a *C. jejuni* infection for a mean of approximately 7.6 years post-infection and observed that these patients were not at increased risk for developing gastrointestinal cancers, but were predisposed to melanomas and squamous cell skin cancers [33]. A significant limitation of this study is that the average time to follow-up is shorter than conservative estimates for the progression of adenomas to carcinomas (5-10 years), which suggests the authors surveyed study participants too early to capture the true impact of *C. jejuni* infection on gastrointestinal cancer development [34]. More recent studies have examined the microbiota of colonic polyps and healthy marginal tissues (HMT) using 16S rRNA gene sequencing, finding that *Campylobacter* genus sequences are enriched in colonic polyps of Italian, Canadian, Chinese, and Spanish cohorts when compared to the HMT [35,36,37]. One study has provided molecular evidence for a link between *C. jejuni* and colorectal cancer by infecting germ-free (GF) Apc^Min/+^mice with both wild-type *C. jejuni* and a CDT mutant, followed by administration of 1% dextran sulfate sodium for 10 days. They found that infection with wild-type *C. jejuni* resulted in increased formation and size of tumors in the distal colon compared to both uninfected mice and mice infected with a CDT mutant, indicating that *C. jejuni* may promote colorectal tumors through the action of CDT. This group also demonstrated that DNA damage was increased in enteroids infected with wild-type *C. jejuni* compared to those infected with the CDT mutant, providing a potential explanation for the mechanism by which *C. jejuni* CDT causes tumorigenesis [38]. Although some of these studies suggest there is an association between *C. jejuni* infection and colorectal cancer, there is still very little information as to whether this is truly causative and what the mechanism of tumorigenesis might be following infection in humans, which underscores the need for more studies in this area.

Important targets to investigate early in the development of colon cancer in response to bacterial infection are hypoxia inducible factor-1 (HIF-1) and activation of the p65 (also called relA) subunit of the NF-κB family of transcription factors by phosphorylation, as both facilitate transcription of hundreds of protumorigenic and cancer progression genes [39]. Hypoxia inducible factor-1 (HIF-1) is a transcriptional regulator which results in the upregulation of hundreds of genes, including those involved in angiogenesis, cell survival and proliferation, metabolism, and tumor metastasis. As a result, elevated HIF-1 is often observed in cancer cells. The HIF-1α subunit is rapidly degraded under normal oxygen conditions, but under hypoxic conditions, HIF-1α is stabilized and leads to HIF-1 promoting expression of its targets [40]. Because hypoxia often occurs as a result of inflammation during bacterial infection, increased HIF-1 activation occurs in host tissues, as well as inflammation through NF-κB. The NF-κB family of transcription factors consists of five subunits, including p65 (also called RelA), whose activation by phosphorylation results in the upregulation of many proinflammatory and protumorigenic targets, including genes involved in cell survival, cell proliferation, angiogenesis, tumor metabolism, and tumor metastasis [41, 42]. Like HIF-1, NF-κB is activated during tumor development and progression and is often elevated in cancer cells [43]. Hypoxia and inflammation caused by bacterial infection could result in tumorigenesis in host tissues via the actions of HIF-1 and NF-κB, a process which, to our knowledge, has yet to be studied in the context of *C. jejuni* infection [44].

In this study, we demonstrate that primary human neutrophils differentiate into discrete subtypes following interaction with *C. jejuni* in a dose and time-dependent manner, including a majority that differentiated into the CD16^hi^/CD62L^lo^ subtype. We visually determined that a significant number of primary human neutrophils exhibited nuclear hypersegmentation during interaction with *C. jejuni*. These neutrophil populations were also found to exhibit reduced apoptosis when compared to uninfected neutrophils. Because the hypersegmented, CD16^hi^/CD62L^lo^ neutrophil subtype has been suggested to inhibit T cells through increased production of arginase-1 and ROS, we quantified these molecules following interaction of *C. jejuni* with primary human neutrophils and found both were significantly elevated [15,5,14]. Such a result suggests that following interaction with *C. jejuni*, neutrophils may contribute to the creation of a protumorigenic environment through inhibition of T cells. To determine the impact of hypersegmented, CD16^hi^/CD62L^lo^ neutrophils on T cells, we measured TCRζ expression in Jurkat T cells after coincubation with *C. jejuni-*infected neutrophils and observed significantly reduced expression of TCRζ. To determine whether arginase-1 or ROS from *C. jejuni-*infected neutrophils was impacting TCRζ expression, we supplemented the media with either L-arginine to counteract the action of arginase-1 or L-ascorbic acid to counteract the action of ROS. We found that supplementation with L-arginine but not L-ascorbic acid restored TCRζ in Jurkats incubated with *C. jejuni-* infected neutrophils. Lastly, because CD16^hi^/CD62L^lo^ neutrophils may impact human colonocytes at the site of infection, we similarly incubated human HCT-116 colonocytes with *C. jejuni*-induced CD16^hi^/CD62L^lo^ neutrophils and observed significantly increased stabilization of the HIF-1α subunit and phosphorylation of the p65 subunit of NF-κB in colonocytes [42]. Taken together, these results suggest that neutrophil subtype differentiation in response to *C. jejuni* infection could facilitate protumorigenic conditions within colonocytes while simultaneously inhibiting the T cell response, allowing tumorigenesis to begin and go undetected by the human immune system. This study is important in progressing our understanding of neutrophil subtypes and *C. jejuni*’s association with colorectal cancer, both of which are understudied and relatively new fields.

### Materials & Methods

1. **Strains and culture conditions:** *C. jejuni* strain 81-176 was grown for 48 hours at 37℃ in microaerobic conditions (85% N_2_, 10% CO_2_, and 5% O_2_) on Mueller-Hinton agar supplemented with 10% sheep’s blood and 10 μg/mL trimethoprim. *Salmonella enterica* serovar Typhimurium strain SL1344 was grown for 24 hours at 37℃ in microaerobic conditions on Mueller-Hinton agar supplemented with 10% sheep’s blood. *Helicobacter pylori* strain 60190 was grown for 72 hours at 37℃ in microaerobic conditions on Mueller Hinton agar supplemented with 10% sheep’s blood.
2. **Isolation of primary human neutrophils:** Human neutrophils were isolated from venous blood from healthy adult volunteers as described previously and in accordance with the Institutional Review Board at the University of Tennessee (UTK IRB-18-04604-XP) [45]. Venous blood was drawn into EDTA-coated vacutainers before being added to 20 mL 1 X phosphate buffered saline (PBS). Then, 10 mL lymphocyte separation medium was underlaid before centrifugation at 1400 rpm for 30 minutes with the brake off. Everything but the red blood cell and neutrophil pellet was aspirated off, and the pellet was resuspended in 20 mL Hank’s balanced salt solution and 20 mL 3% dextran in 0.9% sodium chloride. After a 20-minute incubation at room temperature, the upper layer containing neutrophils was added to a clean tube and centrifuged at 400 x g for 5 minutes. The supernatant was decanted, and the cell pellet washed by resuspending in 20 mL 0.2% sodium chloride, then 20 mL 1.6% sodium chloride was added before centrifuging again at 400 x g for 5 minutes. The supernatant was decanted, and the cell pellet was washed again as described in the previous step. After centrifugation at 400 x g for 5 minutes, the cell pellet was resuspended in RPMI + 10% fetal bovine serum (FBS) and neutrophils were enumerated, and viability was determined using Trypan blue.
3. **Infection of neutrophils with bacteria:** Bacterial cells were suspended in RPMI 1640 + 10% FBS and normalized to an OD600=1.0 before being serially diluted and plated for enumeration of colony forming units (CFU). Unless otherwise specified, neutrophils were infected with the bacterial strains mentioned above in 200 μL RPMI 1640 + 10% FBS at a multiplicity of infection (MOI) of 1:1 (one neutrophil to one bacterial cell) under microaerobic conditions at 37℃.
4. **Flow cytometry analysis of neutrophil populations:** Neutrophils were prepared for flow cytometry as previously described [45]. Unless specified otherwise, 10^6^ neutrophils were used for each replicate. After infection, neutrophils were centrifuged at 400 x g for 5 minutes, washed three times in 1 X PBS, then incubated with Live/Dead Near-IR (NIR) stain (Thermo). Cells were then blocked in SuperBlock blocking buffer in PBS (Thermo) before incubation with antibodies against CD16 and CD62L (Biolegend). Cells were then fixed in fixation buffer (Biolegend) and stored in FACS buffer until analysis. Data was processed using BD and FlowJo software.
5. **Fluorescent microscopy of neutrophil nuclear morphology:** Fluorescent microscopy was performed to assess nuclear morphology. Briefly, 10^6^ neutrophils were added to 35 mm poly-D-lysine coated coverslip dishes (MatTek) in a total volume of 500 μL RPMI 1640 + 10% FBS and incubated under microaerobic conditions at 37℃ for one hour before 10^6^ *C. jejuni* cells were added in 100 μL RPMI 1640 + 10% FBS to a final MOI of 1:1. After incubation for 0, 1, 5, and 24 hours under microaerobic conditions at 37℃, the cells were fixed in fixation buffer (Biolegend), washed three times in 1 X PBS, permeabilized with 0.5% Triton X-100, washed again in 1 X PBS, incubated in SuperBlock blocking buffer in PBS (Thermo) for 30 minutes in microaerobic conditions at 37℃, incubated in 5 μM SYTOX Green in blocking buffer (Invitrogen) for one hour in microaerobic conditions at 37℃, washed in 1 X PBS, and secured with a drop of Mowiol (Sigma) mounting medium. Coverslips were stored in the dark at 4℃ until they were imaged using both fluorescent and brightfield channels. At least 150 individual neutrophils were evaluated for their nuclear morphology per coverslip and timepoint.
6. **Detection of apoptotic neutrophils:** After infection, neutrophils were collected by centrifugation and washed three times with 1 X PBS before being resuspended in 150 μL Annexin binding buffer (Invitrogen). Five μL of Annexin-V AlexaFluor 488 (Invitrogen) was added to each sample and incubated at room temperature for 15 minutes in the dark. Neutrophils were then fixed in fixation buffer (Biolegend) and stored in the dark at 4℃ until flow cytometry analysis. Data was processed using BD and FlowJo software.
7. **Measuring HIF-1α and NF-κB in colonocytes following coincubation with infected neutrophils:** HCT-116 colonic epithelial cells were regularly maintained in DMEM + 10% FBS + 1% L-glutamine + 1% Pen-strep in 5% CO_2_ at 37℃. HCT-116s were seeded overnight in a 24-well tissue culture plate at a density of 10^6^ cells per well in 1 mL DMEM + 10% FBS + 1% L-glutamine + 1% Pen-strep. Before coincubation with neutrophils, the cells were washed in 1 X PBS and 200 μL RPMI 1640 + 10% FBS were added. Subsequently, *C. jejuni* alone, 100 μM cobalt (II) nitrate, or 10^5^ neutrophils which had been uninfected or infected with *C. jejuni* for 5 hours at an MOI of 1:1, were added to the HCT-116s and incubated for one hour in 5% CO_2_ at 37℃. After one hour, the supernatants were removed from the adherent HCT-116s, thoroughly washed three times with 1 X PBS, and then HCT-116 proteins were collected and analyzed by western blot as described below.
8. **Western blot analysis of neutrophil and colonocyte proteins:** After incubation alone or with *C. jejuni*, 10^6^ neutrophils were pelleted at 400 x g and collected in equal volumes 1 X RIPA cell lysis buffer and 2 X Laemmli sample buffer. 10^6^ HCT-116s were collected in 1 X RIPA cell lysis buffer and 2 X Laemmli sample buffer after thorough washing as described above. Proteins were separated on 10% SDS-PAGE gels and transferred at 250 milliamps for 2 hours onto nitrocellulose membranes using transfer buffer consisting of glycine, SDS, and Tris base. The membranes were blocked in 5% nonfat dry milk in TBS-T, then incubated in one of the following antibodies (diluted 1:1000): anti-arginase-1, anti-HIF-1α, anti-p65, anti-phosphorylated p65, or anti-β-actin (Cell Signaling Technology) before detection with the appropriate HRP-conjugated secondary antibody (diluted 1:2000) and Pico-Western luminol/enhancer solution. Densitometry data for western blots were calculated using ImageJ software and normalized to β-actin.
9. **Quantification of reactive oxygen species from neutrophil populations:** After infecting with *C. jejuni* for 5 or 24 hours, 200 μL 1 mM nitroblue tetrazolium chloride was added to 10^6^ primary human neutrophils and incubated for 1 hour at 37℃ under microaerobic conditions. Cells were then collected by centrifugation, resuspended, and lysed in 200 μL 2 M potassium hydroxide, and then solubilized in 200 μL dimethyl sulfoxide. 100 μL of the resulting solution was then added to a 96 well plate and the absorbance was measured at 620 nm [46, 47].
10. **Measuring T cell receptor ζ-chain expression following coincubation with infected neutrophils:** Jurkat T cells were regularly maintained in RPMI 1640 + 10% FBS + 1% L-glutamine + 1% Pen-strep in 5% CO_2_ at 37℃. Before coincubation, 10^6^ Jurkats were pre-incubated in 200 μL RPMI 1640 + 10% FBS that was unsupplemented or supplemented with 1.5 mM L-arginine, 30 μM L-ascorbic acid, or 1 mM L-ascorbic acid. These Jurkats were then incubated overnight with either *C. jejuni* alone or with 10^6^ neutrophils that had either been uninfected or infected with *C. jejuni* for 5 hours in 200 μL RPMI 1640 + 10% FBS supplemented with 1.5 mM L-arginine, 30 μM L-ascorbic acid, or 1 mM L-ascorbic acid. Cells were then analyzed via flow cytometry as described above using antibodies against CD247 (TCRζ) and CD3 (CD3ε) (Biolegend). Data was processed using BD and FlowJo software.
11. **Statistical analysis:** Unless otherwise stated, data were analyzed by one-way ANOVA. Flow cytometry data of CD16 and CD62L cell surface marker expression were analyzed by two-way ANOVA. Arginase-1 western blot data was analyzed by Mann-Whitney T-test. Statistical analysis was performed using Prism 7 software.

## Results

### CD16^hi^/CD62L^lo^ subtype differentiation of *Campylobacter jejuni*-infected neutrophils is MOI- and time-dependent

Neutrophil subtypes are often characterized by CD16 and CD62L cell surface marker expression and nuclear morphology. Neutrophils which highly express both CD16 and CD62L are most commonly isolated from the bloodstream and are associated with the archetypal 3 to 4 lobed nuclei. Neutrophils that express low levels of CD16 but high levels of CD62L are associated with immature neutrophils with banded nuclei. Apoptotic neutrophils often display low levels of CD16 and CD62L [6]. Neutrophils which express high levels of CD16 and low levels of CD62L display hypersegmented nuclei and have been shown to have immunosuppressive effects in other systems, like in sepsis and cancer [9, 10]. As persistent *Campylobacter* infections and reinfection with *Campylobacter* is observed, we suspect that the adaptive immune system may be suppressed in *Campylobacter* infection; however, very little is known about this [21, 22]. Furthermore, hypersegmented, CD16^hi^/CD62L^lo^ neutrophils have also been observed in *Helicobacter pylori* infection, a close relative of *Campylobacter jejuni* which can also present as a persistent infection; however, it has not been elucidated whether hypersegmented, CD16^hi^/CD62L^lo^ neutrophils as a result of *H. pylori* infection have immunomodulatory effects [12].

To determine whether subtypes are observed during *C. jejuni* infection of primary human neutrophils, we used flow cytometry to characterize the neutrophil populations based on the abundance of CD16 and CD62L on individual cells. Initially, we used various multiplicity of infections (MOIs), including 50:1 (bacterial cells-to-neutrophils), 20:1, 5:1, 1:1, and 1:5 (Figure 1 A). After a five-hour incubation, we observed that the majority (61.5%) of uninfected neutrophils displayed CD16^hi^/CD62L^hi^ cell surface marker expression, which are archetypal neutrophils. At 50:1, the majority (65.6%) of neutrophils displayed CD16^lo^/CD62L^hi^ cell surface markers, whereas at 20:1, 48.3% of neutrophils were CD16^lo^/CD62L^hi^ and 48.1% of neutrophils were CD16^lo^/CD62L^lo^. In contrast, at an MOI of 5:1, only 9.5% of neutrophils were CD16^lo^/CD62L^hi^, 39.2% were CD16^lo^/CD62L^lo^, and 45.6% were CD16^hi^/CD62L^lo^. At 1:1, 6.5% of neutrophils were CD16^lo^/CD62L^hi^, and the proportion of CD16^lo^/CD62L^lo^ neutrophils decreased to 24.3% and CD16^hi^/CD62L^lo^ neutrophils increased to 64.5%. At an MOI of 1:5, 7% of neutrophils were CD16^lo^/CD62L^hi^, 18.9% were CD16^lo^/CD62L^lo^, and 70.5% of neutrophils were CD16^hi^/CD62L^lo^. This shows a trend that lower MOIs result in the largest population of CD16^hi^/CD62L^lo^ neutrophils. Overall, these data indicate that the neutrophil response to *C. jejuni* is dependent on bacterial concentrations, which lends remarkable evidence to the plasticity of neutrophils, a concept only recently demonstrated. In addition, since *C. jejuni* infection can be established at low doses, this MOI is likely physiologically relevant to *C. jejuni* infection [48]. Because an MOI of 1:1 induced differentiation of a majority of the neutrophil population to the CD16^hi^/CD62L^lo^ phenotype, the remainder of the experiments were performed at an MOI of 1:1.

**Figure 1:**
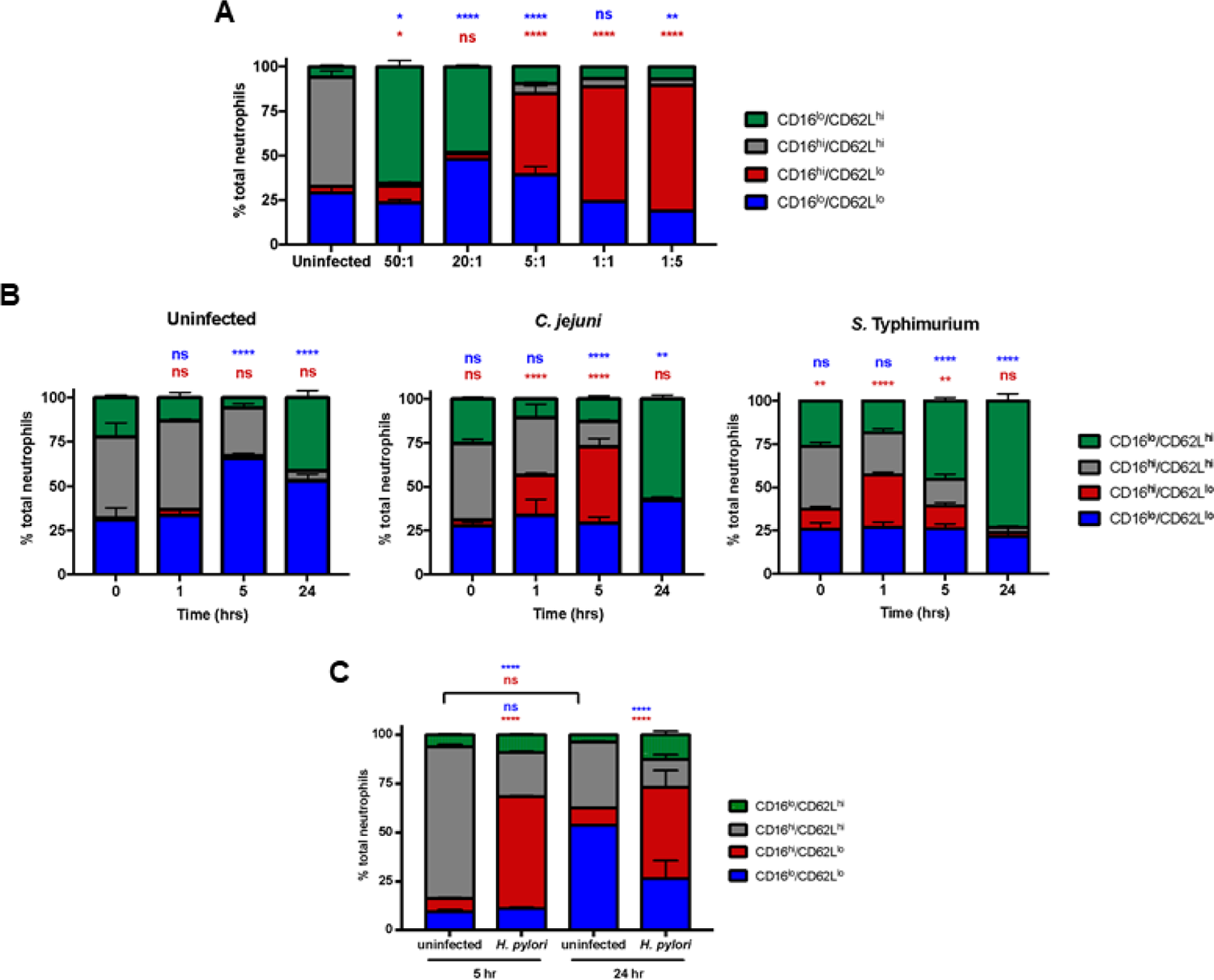
*C. jejuni* induction of CD16^hi^/CD62L^lo^ neutrophils is multiplicity of infection (MOI)- and time-dependent. A. Neutrophils were left uninfected or infected with *C. jejuni* at the indicated varying MOIs for 5 hours. Expression of CD16 and CD62L was measured after 5 hours. B. Neutrophils were left uninfected or infected with *C. jejuni* or *Salmonella* Typhimurium at an MOI of 1:1. Expression of CD16 and CD62L was measured at 0, 1, 5, and 24 hours. C. Neutrophils were left uninfected or infected with *H. pylori* at an MOI of 1:1. Expression of CD16 and CD62L was measured at 5 and 24 hours. Representative graphs, n=3. Percentage of neutrophils are displayed as mean + SEM. Two-way ANOVA. *p<0.05, **p<0.01, ***p< 0.001, ****p<0.0001 compared to A) uninfected, B) 0 hour uninfected, and C) uninfected at corresponding timepoints.

To determine how time impacts differentiation to the CD16^hi^/CD62L^lo^ subtype, neutrophils were infected with *C. jejuni* at an MOI of 1:1 for 0, 1, 5, and 24 hours (Figure 1 B). Also, to compare the neutrophil response toward *C. jejuni* to that of another gastrointestinal pathogen that is also capable of inducing NETs, neutrophils were infected with *Salmonella* Typhimurium strain SL1344 and the CD16 and CD62L markers were examined using the same time points [49]. CD16^hi^/CD62L^lo^ neutrophils were 8.1% of the uninfected population at the zero-hour timepoint, 11.8% of the *C. jejuni-*infected neutrophil population, and was significantly increased for *S.* Typhimurium-infected neutrophils with 26.7% exhibiting the CD16^hi^/CD62L^lo^ subtype. At one hour, 5.7% of uninfected neutrophils were CD16^hi^/CD62L^lo^, 23.1% of *C. jejuni-infected* neutrophils were CD16^hi^/CD62L^lo^, and 30.6% of *S.* Typhimurium-infected neutrophils were CD16^hi^/CD62L^lo^. The results for both *C. jejuni*- and *S.* Typhimurium-infected neutrophils were significantly different at the one-hour mark. By five hours, 5.2% of uninfected neutrophils were CD16^hi^/CD62L^lo^ whereas both *C. jejuni*- and *S.* Typhimurium-infected neutrophils exhibited a significant increase in the CD16^hi^/CD62L^lo^ population at 35.5% and 14.3%, respectively. Although the abundance of CD16^hi^/CD62L^lo^ neutrophils was still significantly increased for *S.* Typhimurium-infected neutrophils at five hours when compared to the zero-hour time point, the population of CD16^hi^/CD62L^lo^ neutrophils decreased at five hours when compared to one hour. By 24 hours of infection with either *C. jejuni* or *S.* Typhimurium, very little of the neutrophil population exhibited the CD16^hi^/CD62L^lo^ phenotype with 1.9% of uninfected neutrophils, 0.4% of *C. jejuni-infected* neutrophils, and 1.2% of *S.* Typhimurium-infected neutrophils presenting as CD16^hi^/CD62L^lo^.

To test this assay using a bacterial strain that has already been found to induce CD16^hi^/CD62L^lo^ neutrophil differentiation, *H. pylori* 60190 was used to infect primary human neutrophils and these cells were similarly analyzed by flow cytometry (Figure 1 C). We observed that *H. pylori*-infected neutrophils exhibited a significant increase in CD16^hi^/CD62L^lo^ neutrophils at five hours (57.4%) and 24 hours (46.8%) when compared to uninfected neutrophils at the same time points (6.8% and 8.9%, respectively). These results confirm those of previous studies and support the methodology used to generate the observations that both *C. jejuni* and *S.* Typhimurium infection lead to significant differences in CD16^hi^/CD62L^lo^ neutrophil populations, albeit for less time, since this subtype is absent by 24 hours post-infection.

### *C. jejuni-*infected neutrophil populations exhibit higher rates of hypersegmented nuclear morphology

Previous studies demonstrated that CD16^hi^/CD62L^lo^ neutrophils possess hypersegmented nuclei, with the number of nuclear lobes often exceeding four [6]. To determine whether the subtype differentiation observed above also leads to changes in nuclear morphology of *C. jejuni*-infected neutrophils, we stained the neutrophil nuclei with Sytox and examined them by fluorescent microscopy (Figure 2). The incidence of hypersegmentation of the neutrophil nucleus, defined as five or greater nuclear lobes, was determined for uninfected and *C. jejuni*-infected neutrophils at 0,1,5, and 24 hours post-infection (Figure 3 A). At zero hours, 6.4% of uninfected neutrophils and 8.0% of *C. jejuni-*infected neutrophils exhibited hypersegmented nuclei. At one hour, 5.3% of uninfected neutrophils and 7.1% of *C. jejuni*-infected neutrophils had hypersegmented nuclei. The incidence of hypersegmented nuclei in *C. jejuni-*infected neutrophils was not significantly different when compared to uninfected neutrophils until the five hour time point, which saw an increase in the incidence of hypersegmentation in *C. jejuni-*infected neutrophils (20.3%) over uninfected (3.1%). At 24 hours, a significant increase in the percentage of hypersegmented nuclei was also observed in *C. jejuni-* infected neutrophils (10.9%) when compared to uninfected (1.5%) cells. Importantly, most of the uninfected neutrophils at 24 hours had condensed, unlobed nuclei (an average of 1.1 lobes per neutrophil), which is indicative of apoptosis; however, *C. jejuni*-infected neutrophil populations exhibited a significantly increased incidence of hypersegmentation at 24 hours, as well as significantly more nuclear lobes on average (2.7 lobes per neutrophil), which indicates prolonged viability (Supplemental Figure 1).

**Figure 2:**
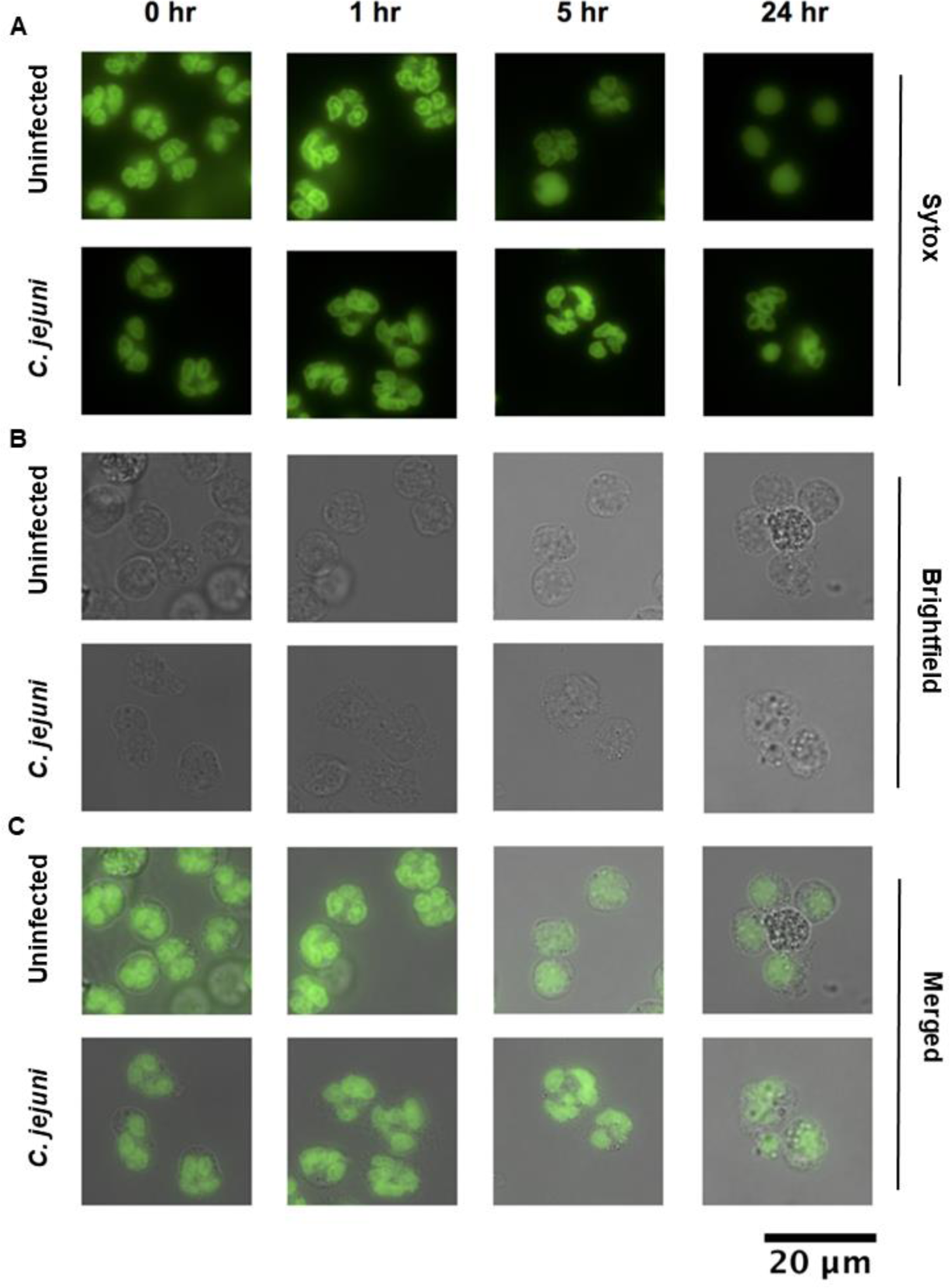
Morphology of neutrophil nuclei upon *C. jejuni* infection. A. Green fluorescence, DNA stained with Sytox. B. Brightfield. C. Merge. Hypersegmentation is defined as >4 nuclear lobes. Condensed, 1 lobed nuclei are indicative of apoptosis. 63X. Representative images, n=3 replicates, >150 neutrophils scored per timepoint.

**Figure 3:**
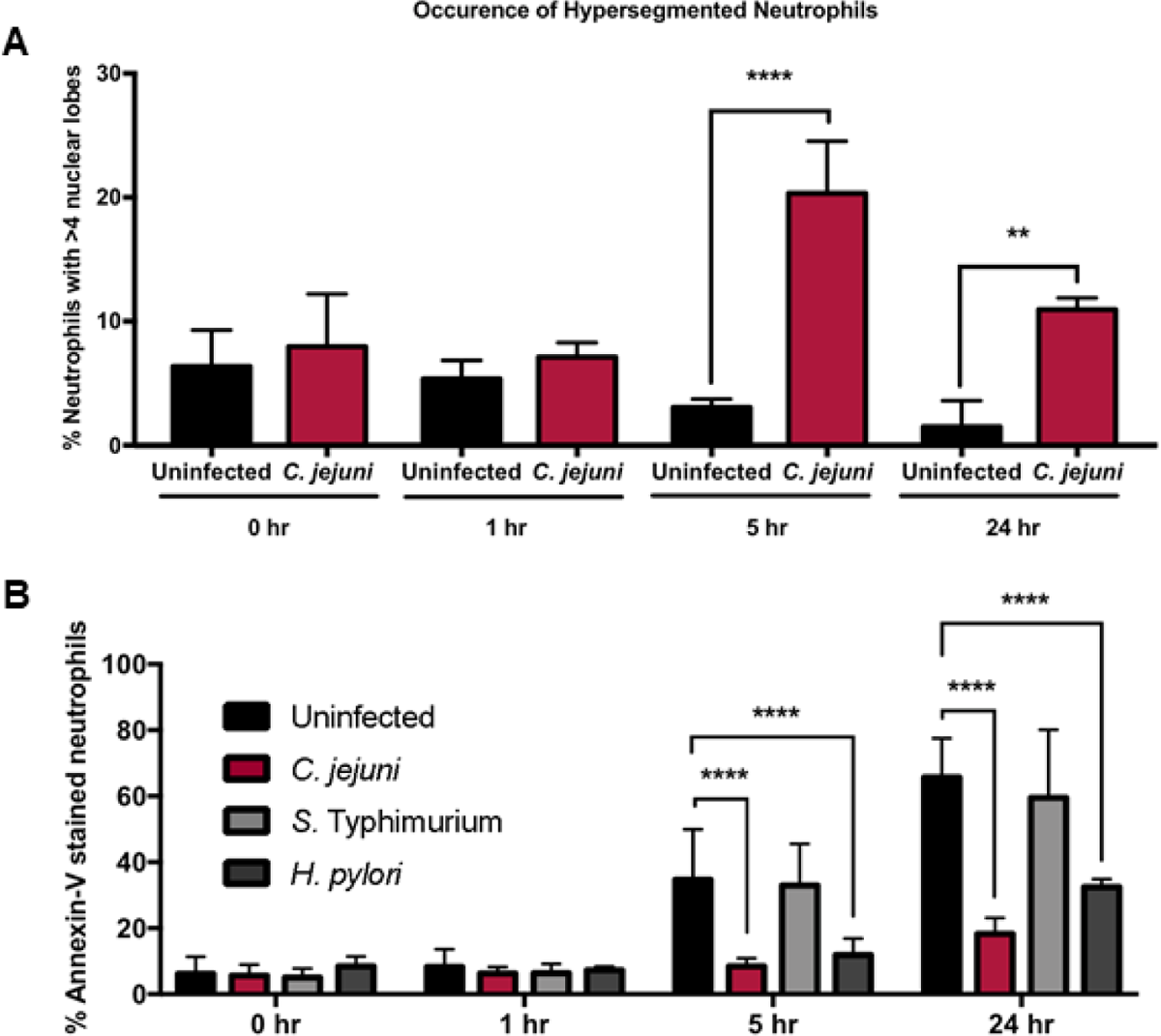
Incidence of hypersegmentation increases and apoptosis decreases at 5 and 24 hours in *C. jejuni*-infected neutrophils. A. Occurrence of hypersegmentation of neutrophil nuclei as measured at 0, 1, 5, and 24 hours in uninfected and *C. jejuni-*infected neutrophils (MOI 1:1). Hypersegmentation is defined as >4 nuclear lobes. B. Occurrence of apoptosis as measured by Annexin-V staining at 0, 1, 5, and 24 hours in uninfected, *C. jejuni*-, *S.* Typhimurium-, and *H. pylori*-infected neutrophils (MOI 1:1). Percentage of neutrophils are displayed as mean + SEM. One-way ANOVA. *p<0.05, **p<0.01, ***p< 0.001, ****p<0.0001 compared to uninfected at corresponding timepoints. A) n=3 replicates, >150 neutrophils scored per timepoint, and B) n=3.

When this microscopy data was compared to the expression of CD16 and CD62L in similar neutrophil populations, a discrepancy emerged at 24 hours since nuclear hypersegmentation was present for *C. jejuni-*infected neutrophils despite only 0.4% of the neutrophil population exhibiting the CD16^hi^/CD62L^lo^ phenotype. We hypothesize that shedding of the CD16 marker occurs between five- and 24-hours post-infection with *C. jejuni* without altering the appearance of the nuclei or the other properties described below; however, further studies should be done to investigate the mechanisms behind this and the differences in CD16 expression at 24 hours during *C. jejuni* and *H. pylori* infection.

### *C. jejuni*-infected neutrophils exhibit reduced apoptosis

Because delayed apoptosis was observed for hypersegmented neutrophils that resulted from infection with *H. pylori* [12], we examined whether *C. jejuni-*infected neutrophils exhibited reduced apoptosis at 0, 1, 5, and 24 hours post-infection (Figure 3 B). No significant changes in apoptosis were observed at the 0 and 1 hour timepoints. At zero hours, 6.2% of uninfected neutrophils were apoptotic, 5.6% of *C. jejuni*-infected, 5.1% of *S.* Typhimurium-infected, and 8.5% of *H. pylori*-infected neutrophils were apoptotic. At one hour, 8.3% of uninfected neutrophils, 6.3% of *C. jejuni*-infected neutrophils, 6.4% of *S.* Typhimurium-infected neutrophils, and 7.3% of *H. pylori-*infected neutrophils were apoptotic. In contrast, apoptosis was significantly decreased at 5 and 24 hours post-infection for neutrophils infected with *C. jejuni* or *H. pylori* when compared to either uninfected or *S.* Typhimurium-infected neutrophils. Specifically, at five hours, 34.7% of uninfected neutrophils were apoptotic whereas 8.5% and 11.9% of *C. jejuni* or *H. pylori-* infected neutrophils, respectively, were apoptotic. By comparison, 32.9% of *S.* Typhimurium-infected neutrophils were apoptotic at five hours, which was not significantly different from uninfected populations. At 24 hours, the majority of uninfected neutrophils (65.7%) and *S.* Typhimurium-infected neutrophils (59.5%) were apoptotic, whereas only 18.3% of *C. jejuni*-infected neutrophils and 32.4% of *H. pylori-* infected neutrophils were apoptotic. These results from *C. jejuni* and *H. pylori-*infected neutrophil populations were significantly different when compared to uninfected or *S.* Typhimurium-infected neutrophil populations. Significantly delayed neutrophil apoptosis upon infection with *C. jejuni* is important because it not only further supports that hypersegmented, CD16^hi^/CD62L^lo^ neutrophils are induced during *C. jejuni* infection, but also that extending the life of neutrophils may be responsible for the immunopathological effects that occur during human *C. jejuni* infection.

### *C. jejuni-*infected neutrophils exhibit increased arginase-1 expression and reactive oxygen species production

Hypersegmented, CD16^hi^/CD62L^lo^ neutrophils have been previously shown to negatively impact T cell activation and proliferation by reducing expression of the ζ-chain of the T cell receptor (TCRζ) through either the production of arginase-1 or the generation of reactive oxygen species [14]. To determine whether the hypersegmented, CD16^hi^/CD62L^lo^ neutrophils induced by *C. jejuni* infection could impact T cells, we first examined the production of arginase-1 and reactive oxygen species within infected neutrophil populations.

To examine arginase-1 production, *C. jejuni* was used to infect primary human neutrophils under the same conditions where a majority of the population exhibits the CD16^hi^/CD62L^lo^ phenotype. As expected, a significant, approximately two-fold increase of arginase-1 was observed for *C. jejuni-*infected neutrophil (density relative to β-actin=0.9) whole cell lysates when compared to those from uninfected neutrophils (density relative to β-actin= 0.4) (Figure 4 A&B). As hypersegmented, CD16^hi^/CD62L^lo^ neutrophils have also been shown to reduce TCRζ expression by elevated production of ROS, we used nitroblue tetrazolium chloride to examine the ability of *C. jejuni*-infected neutrophils to produce ROS. Nitroblue tetrazolium chloride is one of the most common methods to measure the respiratory burst and ROS production in neutrophils. Reduction of nitroblue tetrazolium chloride by ROS forms an indigo-colored, insoluble formazan precipitate, which is then solubilized and whose absorbance is measured [47]. Using the same conditions where a majority of the infected neutrophil population exhibits the CD16^hi^/CD62L^lo^ phenotype, a significant, three-fold increase in ROS production was observed when compared to uninfected neutrophils (Figure 4 C). To further support our hypothesis that the CD16^hi^/CD62L^lo^ neutrophil subtype is specifically responsible for the increase in ROS and not a general response to *C. jejuni* infection of neutrophils, neutrophils were infected with *C. jejuni* at an MOI of 20:1 for 5 hours and ROS was similarly measured. This is due to the above observation that infection of neutrophils with *C. jejuni* for five hours at an MOI of 20:1 results in predominantly CD16^lo^/CD62L^hi^ and CD16^lo^/CD62L^lo^ neutrophil populations. As a result, neutrophils infected at an MOI of 20:1 produced approximately 70% of the ROS produced by neutrophils infected with an MOI of 1:1, which supports other studies demonstrating the unique ability of CD16^hi^/CD62L^lo^ neutrophils to produce elevated amounts of ROS (Supplemental Figure 2).

**Figure 4:**
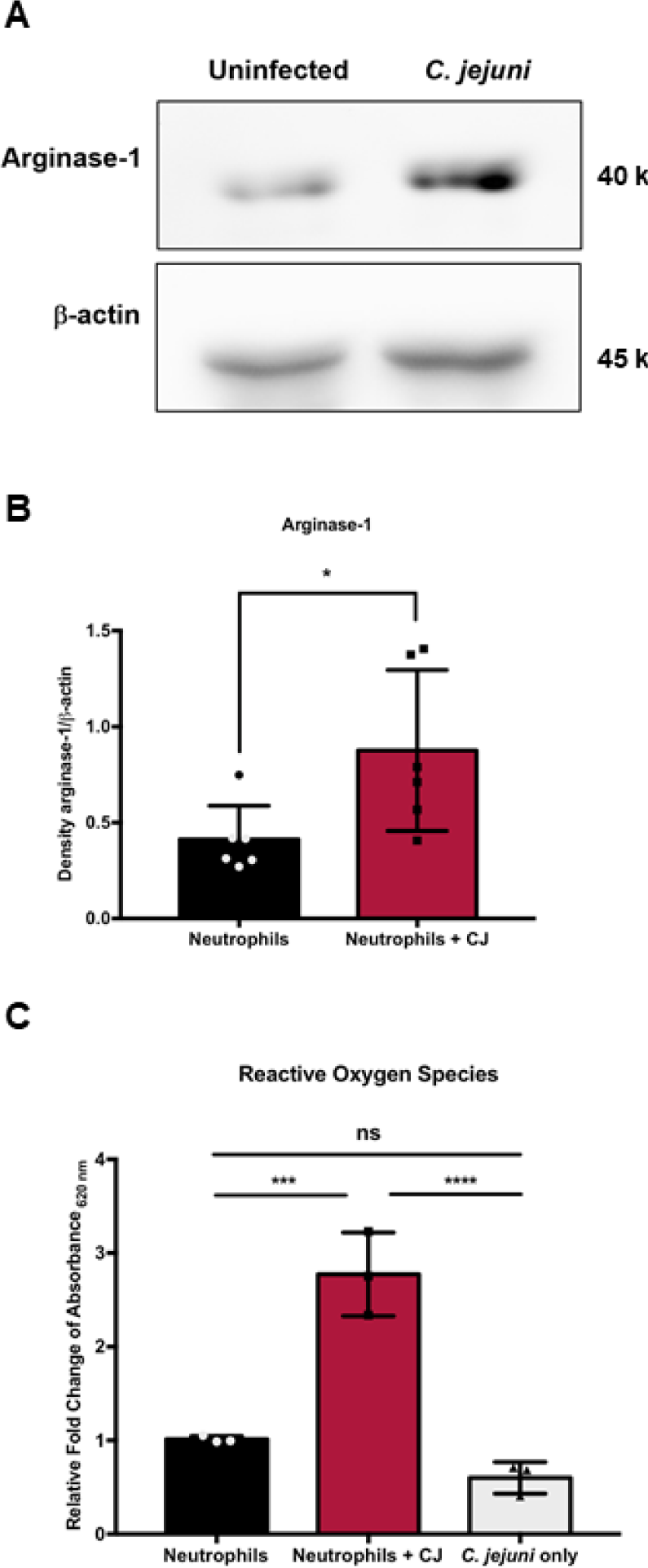
Arginase-1 expression and reactive oxygen species production are increased in neutrophils infected with *C. jejuni* (MOI 1:1) for 5 hours compared to uninfected. A. Western blot of uninfected and *C. jejuni*-infected neutrophil lysates. B. Densitometry of arginase-1 relative to β-actin. C. Relative fold-change compared to uninfected neutrophils from absorbance at 620 nm values. Densitometry and fold change displayed as mean + SEM. B) Mann-Whitney test, and C) One-way ANOVA *p<0.05, **p<0.01, ***p< 0.001, ****p<0.0001 compared to uninfected. A) representative image, n=3, B) n=3, and C) n=3.

### Coincubation of *C. jejuni*-infected neutrophils with human T cells leads to reduced TCRζ expression

To determine whether hypersegmented, CD16^hi^/CD62L^lo^ neutrophils induced by *C. jejuni* infection impact the T cell receptor, we incubated neutrophils with *C. jejuni* under conditions where a majority differentiate to the CD16^hi^/CD62L^lo^ phenotype. We then incubated these or uninduced neutrophils with human T cells overnight and examined for TCRζ expression by flow cytometry (Figure 5, Supplemental Figure 3). From this analysis, we observed that 73.5% of T cells possessed detectable TCRζ following coincubation with *C. jejuni*-infected neutrophils when compared to T cells alone (100%). This is a significant reduction in the TCRζ expressing population, including when T cells were incubated with uninfected human neutrophils (99.0%) or with *C. jejuni* alone (96.0%) (Figure 5 A). To determine whether increased arginase-1 production by *C. jejuni*-infected neutrophils was responsible for the reduction in TCRζ expression, T cells were supplemented with 1.5 mM L-arginine for 24 hours before and during coincubation to counteract the effects of arginase-1. This concentration of L-arginine was chosen as it has been shown that 1 to 2 mM L-arginine maximally activates CD3+ T cells [50, 51]. In the presence of L-arginine, TCRζ expression of T cells incubated with uninfected neutrophils (105.1%), with *C. jejuni-*infected neutrophils (89.5%), or with *C. jejuni* alone (101.1%) was not significantly decreased when compared to T cells alone supplemented with L-arginine (100%) (Figure 5 B).

**Figure 5:**
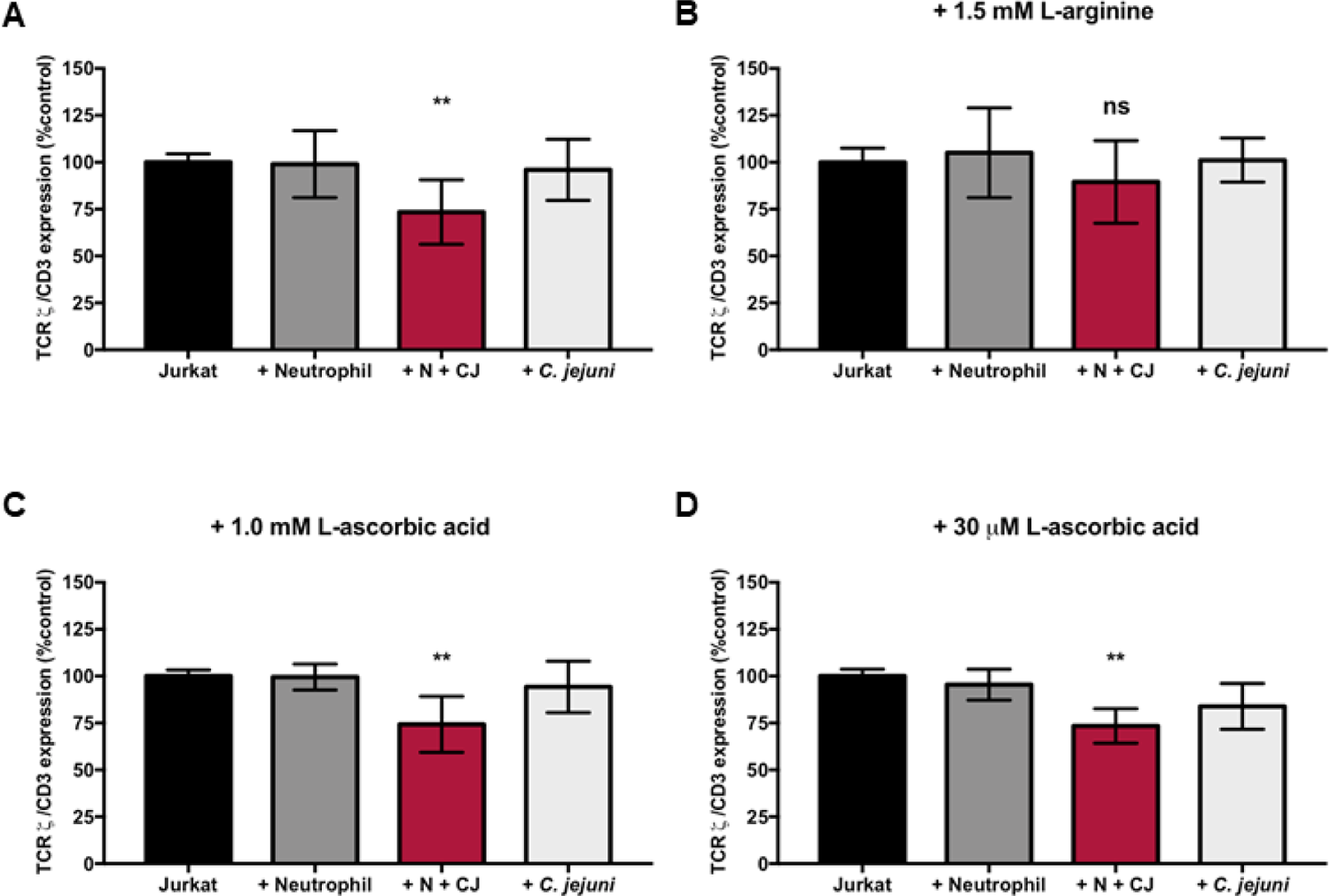
TCRζ chain reduction in Jurkats incubated with *C. jejuni*-infected neutrophils is restored in the presence of 1.5 mM L-arginine, not 1.0 mM or 30 µM L-ascorbic acid. A. TCRζ expression relative to CD3 normalized to Jurkats alone in RPMI 1640 + 10% FBS. B. TCRζ expression relative to CD3 normalized to Jurkats alone in RPMI 1640 + 10% FBS + 1.5 mM L-arginine. C. TCRζ expression relative to CD3 normalized to Jurkats alone in RPMI 1640 + 10% FBS + 1.0 mM L-ascorbic acid. D. TCRζ expression relative to CD3 normalized to Jurkats alone in RPMI 1640 + 10% FBS + 30 µM L-ascorbic acid. %TCRζ expression expressed as mean + SEM. One-way ANOVA *p<0.05, **p<0.01, ***p< 0.001, ****p<0.0001 compared to uninfected A) n=5, B) n=3, C) n=3, and D) n=3.

Because ROS produced by hypersegmented, CD16^hi^/CD62L^lo^ neutrophils has also been shown to inhibit TCRζ expression, T cells were supplemented with either 1 mM or 30 μM L-ascorbic acid, an antioxidant, before and during coincubation to counteract ROS produced by *C. jejuni*-infected neutrophils. These concentrations were chosen, as 1 mM is a supraphysiological concentration of L-ascorbic acid in T cells, whereas 30 μM is a physiologically normal concentration of L-ascorbic acid in T cells and serum [52]. The addition of either concentration of L-ascorbic acid did not affect TCRζ expression when compared to T cells incubated without supplementation.

Specifically, upon addition of 1 mM L-ascorbic acid, we observed no significant changes to the T cell population expressing TCRζ when incubated with uninfected neutrophils (99.5%) or incubated with *C. jejuni* alone (94.3%) when compared to T cells alone (100%). However, like our unsupplemented group, we still observed a significant decrease in TCRζ expression in T cells when incubated with *C. jejuni*-infected neutrophils (74.3%) despite the addition of 1 mM L-ascorbic acid (Figure 5 C). This result was also observed upon supplementation with 30 μM L-ascorbic acid. For example, a significant decrease in the T cell population expressing TCRζ was observed following incubation with *C. jejuni-*infected neutrophils (73.4%) and no significant differences were detected following incubation of T cells with uninfected neutrophils (95.5%) or *C. jejuni* alone (83.9%) when compared to T cells alone supplemented with 30 μM L-ascorbic acid (Figure 5 D).

### Coincubation of *C. jejuni*-infected neutrophils with human colonocytes induces HIF-1α stabilization and phosphorylation of NF-κB

Beyond impacts to T cells, the significantly increased production of ROS and inflammation from hypersegmented, CD16^hi^/CD62L^lo^ neutrophils may also affect adjacent colonocytes. To determine the potential effects of CD16^hi^/CD62L^lo^ neutrophil-derived ROS production on the colonic epithelium, we first examined HIF-1α stabilization in colonocytes. HIF-1 is a transcription factor that promotes the expression of several systems involved in tumorigenesis, and ROS has been shown to increase HIF-1α subunit stabilization [53]. We incubated predominantly CD16^hi^/CD62L^lo^ neutrophil populations or uninfected neutrophils with colonocytes for one hour and examined for HIF-1α stabilization by western blot. In addition, we treated colonocytes with cobalt (II) as a positive control since it has been shown to stabilize HIF-1α independent of hypoxia [54]. Colonocytes incubated with *C. jejuni*-infected neutrophils, which are predominantly the CD16^hi^/CD62L^lo^ subtype, exhibited an approximately two-fold increase in HIF-1α stabilization (density relative to β-actin=1.8), whereas incubation of colonocytes with *C. jejuni* alone (density relative to β-actin=0.90) or uninfected neutrophils (density relative to β-actin=0.95) failed to produce any significant changes in HIF-1α stabilization when compared to colonocytes alone (density relative to β-actin=0.96). In contrast, colonocytes incubated with cobalt (II) nitrate exhibited a 3.5-fold increase in HIF-1α (density relative to β-actin=3.4) (Figure 6 A&B). We hypothesize that increased production of ROS by hypersegmented, CD16^hi^/CD62L^lo^ neutrophils that are induced in response to *C. jejuni* are responsible for increased stabilization of HIF-1α in these colonocytes, which may lead to transcription of pro-tumorigenic genes targets of HIF-1.

**Figure 6:**
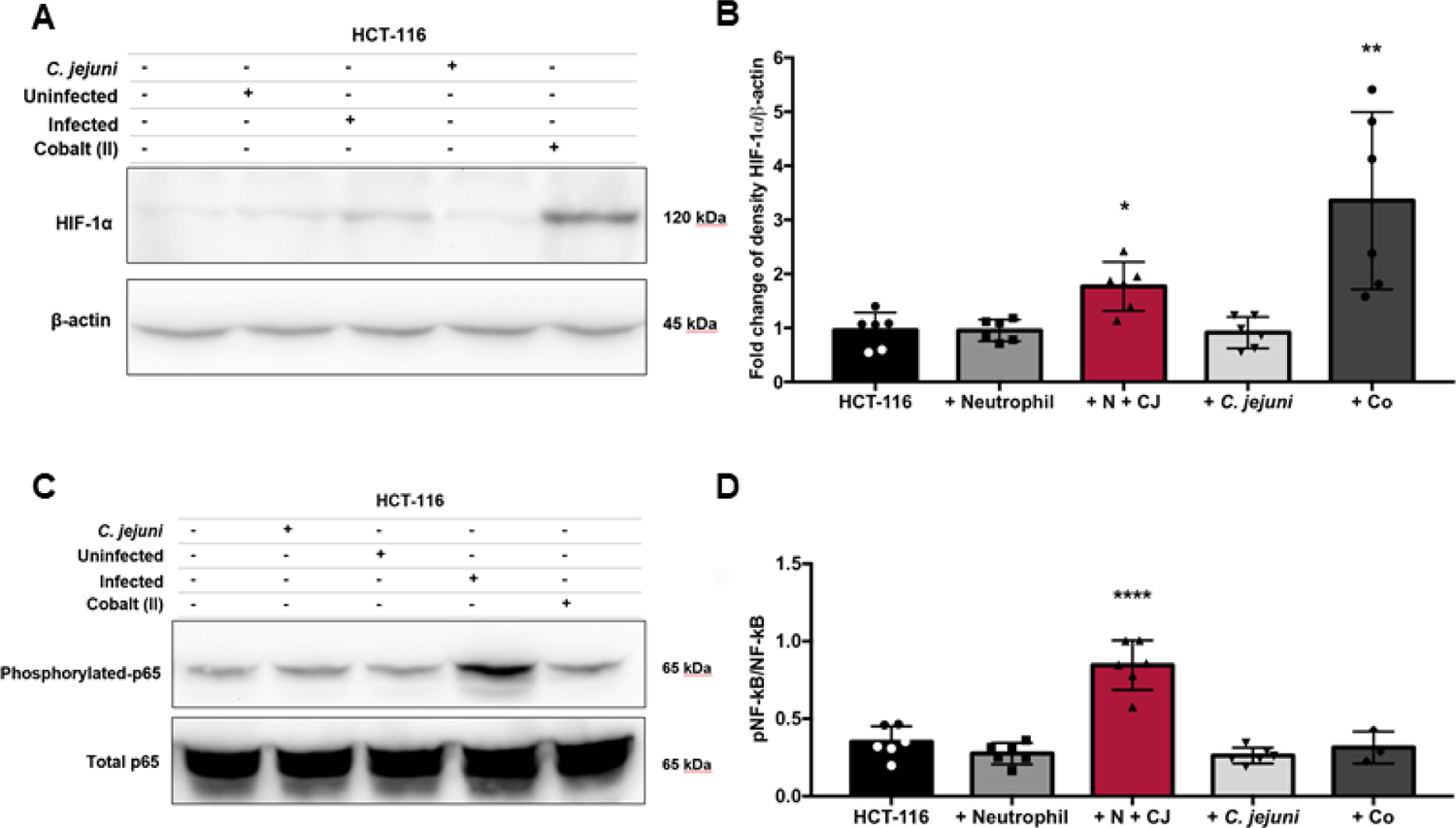
HIF-1a stabilization and p65 phosphorylation of NF-κB are increased in HCT-116 colonocytes coincubated with *C. jejuni-*infected neutrophils. A. Western blot of HCT-116 colonocyte cell lysates alone or after 1-hour coincubation with uninfected neutrophils, neutrophils infected with *C. jejuni* (MOI 1:1) for 5 hours, *C. jejuni* alone, or cobalt (II) nitrate as a positive control. B. Densitometry of HIF-1α relative to β-actin. C. Western blot of HCT-116 colonocyte cell lysates alone or after 1-hour coincubation with uninfected neutrophils, *C. jejuni* alone, neutrophils infected with *C. jejuni* (MOI 1:1) for 5 hours, or cobalt (II) nitrate. D. Densitometry of the phosphorylated p65 subunit of NF-κB relative to the total p65 subunit of NF-κB. Densitometry displayed as mean + SEM. A) nonparametric test, and B) One-way ANOVA *p<0.05, **p<0.01, ***p< 0.001, ****p<0.0001 compared to uninfected. A) representative image, n=3, B) n=3, C) representative image, n=3, and D) n=3.

**Figure 7:**
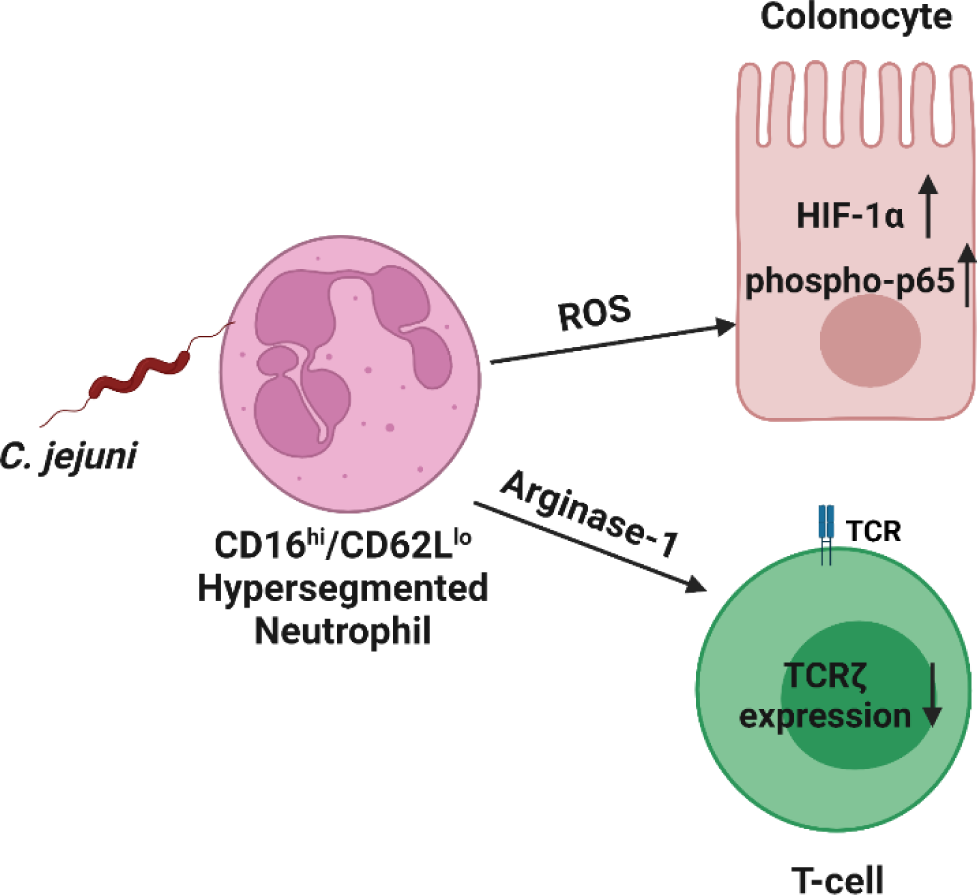
Graphical Abstract: Infection of neutrophils with *C. jejuni* results in the formation of a hypersegmented, CD16^hi^/CD62L^lo^ neutrophil subtype in an MOI- and time-dependent manner. *C. jejuni*-infected neutrophils produce elevated levels of reactive oxygen species and show elevated expression of arginase-1. The increased production of reactive oxygen species may be responsible for increased HIF-1α stabilization and phosphorylation of the p65 subunit of the NF-κB family of transcription factors in colonocytes coincubated with *C. jejuni*-infected neutrophils. The increased expression of arginase-1 may be responsible for the downregulation of the ζ-chain of the T cell receptor when T cells are incubated with *C. jejuni*-infected neutrophils.

HIF-1 and NF-κB have been shown to interact during infection and inflammation, as NF-κB is often activated when HIF-1α is stabilized [55]. Because we have demonstrated that HIF-1α is stabilized in colonocytes incubated with *C. jejuni-*infected neutrophils, we also examined NF-κB activation through phosphorylation of p65 in these colonocytes (Figure 6 C&D). As expected, phosphorylation of p65 was significantly increased (2.4 fold) in colonocytes incubated with neutrophil populations where the CD16^hi^/CD62L^lo^ subtype is the majority. In contrast, colonocytes incubated with uninfected neutrophils or *C. jejuni* alone showed no significant change in phosphorylation of p65. This was somewhat surprising since other groups have demonstrated *C. jejuni* alone induces NF-κB activation in colonocytes [56]. Instead, our results suggest that neutrophil activity, including the induction of the hypersegmented, CD16^hi^/CD62L^lo^ subtype, may be responsible for increased inflammation and resulting pathology of host intestinal tissue. Interestingly, colonocytes incubated with cobalt (II) nitrate showed no significant change in phosphorylation of p65. This suggests that NF-κB activation may occur upstream of HIF-1α stabilization in response to ROS production in this system, as cobalt acts directly on HIF-1α alone.

## Discussion

Infection by *Campylobacter* species is a significant cause of gastrointestinal disease worldwide that has both acute and long-term implications to human health, including the development of post-infectious inflammatory disorders and possibly colorectal cancer [19,20,38,35,36,37]. Importantly, the bacterial and host factors that are responsible for these various outcomes are mostly unknown. For the field to advance in understanding the processes that contribute to *Campylobacter-*induced diseases, it is important that we begin to thoroughly characterize the host responses to these important pathogens and how those processes impact the host. We have demonstrated in this study that the majority of human neutrophils differentiate into a hypersegmented, CD16^hi^/CD62L^lo^ subtype after being exposed to low doses of *C. jejuni* and that the neutrophil population exhibits delayed apoptosis, increased expression of arginase-1, and elevated production of ROS. Not only do these neutrophil activities support our initial observations that *C. jejuni* infection induces differentiation of the hypersegmented, CD16^hi^/CD62L^lo^ subtype, but it also suggests how immunopathology during campylobacteriosis might occur. For example, the decreased apoptosis of neutrophil populations we observed during infection likely leads to prolonged exposure of the surrounding gastrointestinal tissues to the activities of neutrophils [1, 57]. This includes several processes and products we previously examined that may directly damage host cells or are exceedingly proinflammatory, including the release of S100A12, lipocalin-2, myeloperoxidase, and neutrophil elastase [25, 49]. Beyond these more classical responses, our current work suggests that a portion of the neutrophil population may be inappropriately immunosuppressive and counteract specific components of adaptive immunity (e.g., T cells). In addition, it is particularly interesting to note that induction of the hypersegmented, CD16^hi^/CD62L^lo^ neutrophil subtype is greatest when there are one or fewer bacteria per neutrophil, since we previously demonstrated that NET formation occurs when there are ten or greater *C. jejuni* cells to one neutrophil [49]. These results indicate that neutrophils dynamically respond to stimuli at the site of infection, which i) is in stark contrast to early impressions of neutrophils as terminally differentiated and transcriptionally limited cells and ii) suggests that host responses and disease outcomes may be dependent on *Campylobacter* burden.

Further, hypersegmented, CD16^hi^/CD62L^lo^ neutrophils as well as myeloid derived suppressor cells (MDSCs) produce elevated levels of arginase-1 and reactive oxygen species [14]. In order to further classify the hypersegmented, CD16^hi^/CD62L^lo^ neutrophil differentiation we observed in response to *C. jejuni* infection and to investigate what effects the neutrophil subtype may have on the host during campylobacteriosis, we examined arginase-1 expression and reactive oxygen species production. As predicted, we found that both were elevated in neutrophils which had been incubated with *C. jejuni* under conditions where a majority develop into the hypersegmented, CD16^hi^/CD62L^lo^ subtype. The ability of this neutrophil subtype and MDSCs to suppress T cell function has been attributed to decreased T cell receptor ζ-chain (TCRζ) expression, which is caused by either L-arginine depletion via arginase-1 or reactive oxygen species directly impacting the TCR [15,16,18]. Because of this, we subsequently demonstrated that human T cells incubated with these differentiated neutrophil populations exhibited a significant decrease in TCRζ expression, which is important for T cell receptor signaling and proliferation. To preliminarily determine whether arginine depletion or ROS production were responsible for this reduction in TCRζ expression, we supplemented T cells with exogenous L-arginine or an antioxidant (L-ascorbic acid), finding that L-arginine partially restored TCRζ expression in the T cell population while L-ascorbic acid did not. Determining whether arginase-dependent suppression of T cells occurs during *C. jejuni* infection could provide valuable targets for aiding the adaptive immune system’s response to *C. jejuni* infection. This could help reduce persistent infection or reinfection, aid in vaccine development through the elicitation of an adequate memory response or provide protection against cancer formation by strengthening the adaptive immune response [17]. Indeed, dietary supplementation with L-arginine has been shown to increase bacterial clearance and decrease susceptibility to bacterial infections and sepsis [58, 59]. Better nutrition and increased bacterial clearing as a result of dietary L-arginine in developed countries may explain why persistent *Campylobacter* infections occur less frequently in developed countries than in developing countries [60, 19]. The interactions between *C. jejuni* infection and nutrition, especially arginine, and their global trends are areas which need further investigation.

Beyond their impacts to inflammation and T cell activation, hypersegmented, CD16^hi^/CD62L^lo^ neutrophils could have direct implications to colonocytes, including colorectal tumorigenesis. Although the links between *C. jejuni* infection and colorectal cancer is severely understudied, *C. jejuni* has been proposed to cause colorectal tumors through damage to colonocyte DNA by cytolethal distending toxin (CDT) [38]. Furthermore, 16S rRNA gene sequencing studies have shown members of the *Campylobacter* genus are associated with colorectal polyps and tumors but not healthy marginal tissue [35,36,37]. These studies suggest that sample type may have impacted the field’s ability to associate *Campylobacter* infection with colorectal tumorigenesis since an initial study that established a correlation does not exist relied on fecal samples for detection [33]. If *Campylobacter* remains restricted to polyp or tumor tissues, as the tissue microbiome studies suggest, culture or PCR of fecal samples may not be sufficient to detect these associations. Further supporting a possible association with colorectal cancer, we found that incubation of colonocytes with neutrophil populations where hypersegmented, CD16^hi^/CD62L^lo^ cells predominate, HIF-1α stabilization and NF-κB activation occurred, a combination which could lead to upregulation of hundreds of tumor promoting genes. An unexpected observation from this work was that, despite repeated attempts using various cell lines, *C. jejuni* strains, and MOIs, we were unable to observe NF-κB activation in the absence of neutrophils [data not shown]. In the end, damage to colonocytes at the site of infection directly through *C. jejuni* effectors like CDT or through the activities of the innate immune response (e.g. NETosis) combined with stabilization of HIF-1α, activation of NF-κB, and decreased T cell function through TCRζ downregulation, could allow for tumorigenesis to occur and go unchecked. As T cells serve an important role in surveillance of host tissue for cancer, further studies need to be conducted to determine whether immunosuppressive neutrophil subtypes, as well as direct effects of *C. jejuni* on colonocytes, could be responsible for the development of colorectal cancer.

## Conclusions

We have shown that *Campylobacter jejuni*-infected neutrophils will differentiate into a hypersegmented, CD16^hi^/CD62L^lo^ subtype in an MOI- and time-dependent manner. These *C. jejuni*-infected neutrophils display delayed apoptosis and increased arginase-1 and reactive oxygen species production. Coincubation of T cells with *C. jejuni*-infected neutrophils results in a decrease in T cell populations expressing TCRζ, which is restored when L-arginine is added, but not when the antioxidant L-ascorbic acid is added, indicating that L-arginine depletion by arginase-1 may be responsible for decreased TCRζ expression. Further, coincubation of colonocytes with *C. jejuni*-infected neutrophils results in increased stabilization of HIF-1α and increased phosphorylation of the p65 subunit of NF-κB in the colonocyte. When these results are taken together, *C. jejuni*-infected neutrophils may result in the transcription of many pro-tumorigenic genes by HIF-1 and NF-κB in the colonocyte and suppress the ability of T cells to respond to both the infection and cancerous antigens in the affected colonocytes. This study highlights the importance and dynamic nature of neutrophils in response to *C. jejuni* infection and may provide targets for treatment and insights regarding the association of *C. jejuni* infection and cancer.

## Data Availability Statement

All data will be made available upon request to the corresponding author.

## Acknowledgements

The authors would like to thank Trevor J. Hancock for assistance with flow cytometry and experimental advice, Jenny K. Heppert and Sarah J. Kauffman for microscopy assistance and advice, Jaydeep Kolape at the UTK Advanced Microscopy Center, Bohye Park, Rachel Patton McCord, Tim Sparer, and Benjamin J. Parker for advice, and Eleanor Mancini and Caroline Parker for their assistance. Graphical abstract was created with BioRender software.

