## Supplemental for "*Campylobacter jejuni* induces differentiation of human neutrophils to the CD16^hi^/CD62L^lo^ subtype which possess cancer promoting activities"

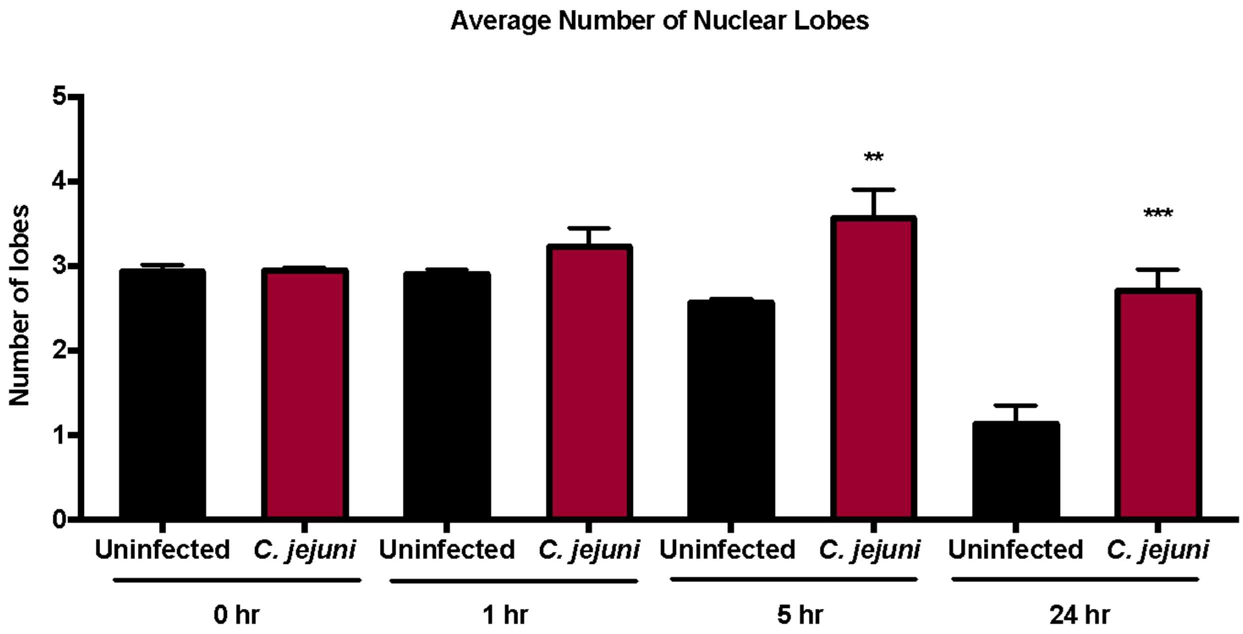


Supplemental Figure 1: Average number of nuclear lobes increases at 5 and 24 hours in *C. jejuni* infected neutrophils.
Number of lobes of each neutrophil nuclei as measured at 0, 1, 5, and 24 hours in uninfected and *C. jejuni* infected neutrophils (MOI 1:1).
One-way ANOVA. *p<0.05, **p<0.01, ***p< 0.001, ****p<0.0001 compared to uninfected at corresponding timepoints. n=3 replicates, >150 neutrophils scored per timepoint.

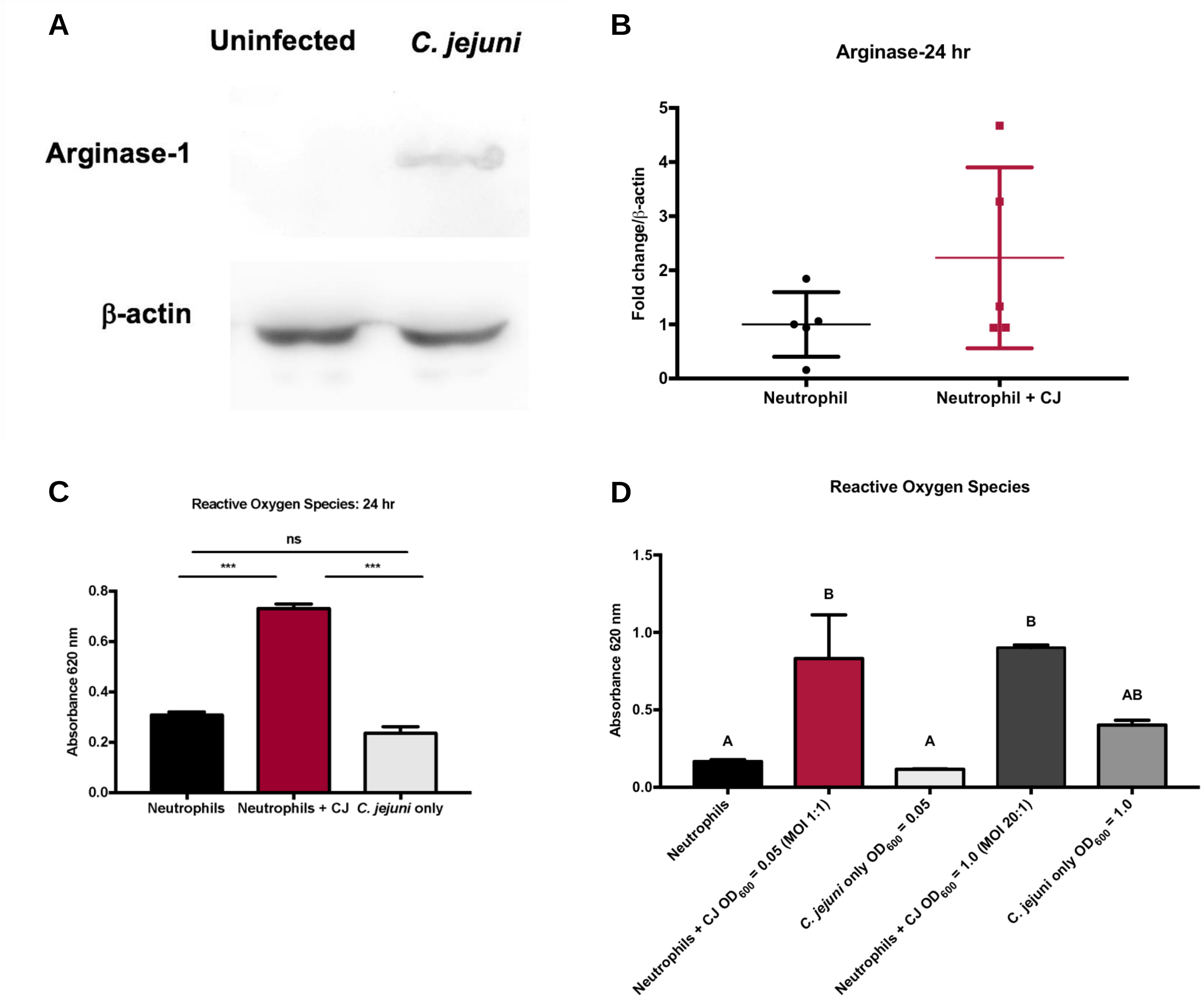


Supplemental Figure 2: Arginase-1 expression and reactive oxygen species production are still increased in neutrophils infected with *C. jejuni* (MOI 1:1) for 24 hours compared to uninfected and reactive oxygen species generation is decreased in neutrophils infected with *C. jejuni* at an MOI of 20:1 compared to neutrophils infected with *C. jejuni* at an MOI of 1:1.

A. Western blot of uninfected and *C. jejuni* infected neutrophil lysates.
B. Densitometry of arginase-1 relative to β-actin.
C. Absorbance at 620 nm values.
D. Absorbance at 620 nm values.

Densitometry and absorbance values displayed as mean + SEM. B) Mann-Whitney test, and B) One-way ANOVA *p<0.05, **p<0.01, ***p< 0.001, ****p<0.0001 compared to uninfected. A) representative image, n=3, B) n=3, C) n=2, and D) n=2.

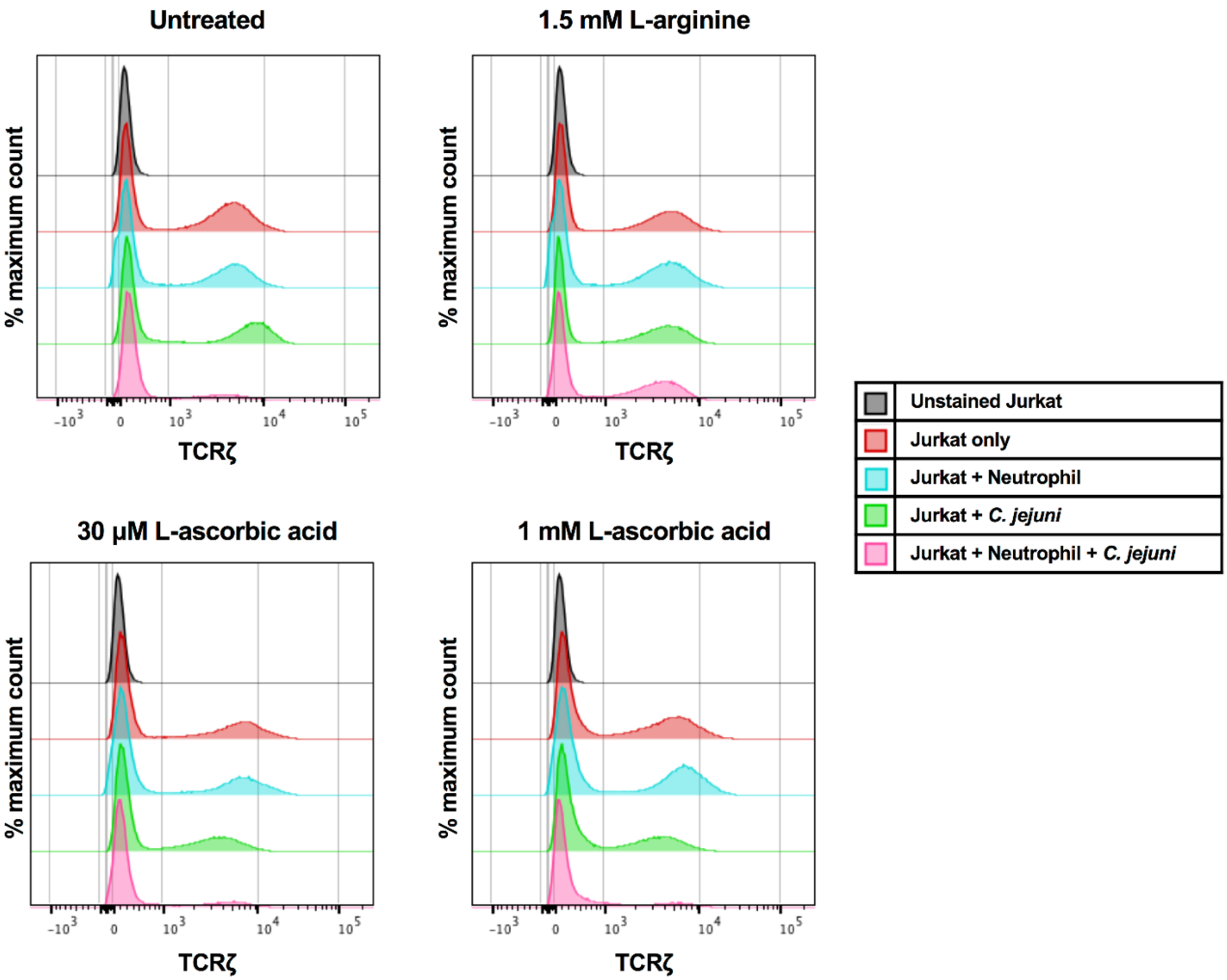


Supplemental Figure 3: TCRζ reduction in Jurkats coincubated with *C. jejuni* infected neutrophils is restored in the presence of 1.5 mM L-arginine, not 1.0 mM or 30 µM L-ascorbic acid.

A. TCRζ expression of unstained Jurkats, Jurkats alone, Jurkats + Neutrophils, Jurkats + Neutrophils + *C. jejuni*, and Jurkats + *C. jejuni* in RPMI 1640 + 10% FBS.
B. TCRζ expression of unstained Jurkats, Jurkats alone, Jurkats + Neutrophils, Jurkats + Neutrophils + *C. jejuni*, and Jurkats + *C. jejuni* in RPMI 1640 + 10% FBS + 1.5 mM L-arginine.
C. TCRζ expression of unstained Jurkats, Jurkats alone, Jurkats + Neutrophils, Jurkats + Neutrophils + *C. jejuni*, and Jurkats + *C. jejuni* in RPMI 1640 + 10% FBS + 30 µM L-ascorbic acid.
D. TCRζ expression of unstained Jurkats, Jurkats alone, Jurkats + Neutrophils, Jurkats + Neutrophils + *C. jejuni*, and Jurkats + *C. jejuni* in RPMI 1640 + 10% FBS + 1.0 mM L-ascorbic acid.

TCRζ expression counts are normalized to mode and expressed as % maximum count A) n=5, B) n=3, C) n=3, and D) n=3. Representative images.
